# 5’ UTR recruitment of eIF4GI or DAP5 drives cap-independent translation for a subset of human mRNAs

**DOI:** 10.1101/472498

**Authors:** Solomon A. Haizel, Usha Bhardwaj, Ruben L. Gonzalez, Somdeb Mitra, Dixie J. Goss

## Abstract

During unfavorable human cellular conditions (*e.g*., tumor hypoxia, viral infection, *etc*.), canonical, cap-dependent mRNA translation is suppressed. Nonetheless, a subset of physiologically important mRNAs (*e.g*., HIF-1α, FGF-9, and p53) is still translated by an unknown, cap-independent mechanism. Additionally, expression levels of eIF4G and its homolog, death associated protein 5 (DAP5), are elevated. Using fluorescence anisotropy binding studies, luciferase reporter-based *in vitro* translation assays, and mutational analyses, here we demonstrate that eIF4GI and DAP5 specifically bind to the 5’ UTRs of these cap-independently translated mRNAs. Surprisingly, we find that the eIF4E binding domain of eIF4GI increases not only the binding affinity, but also the selectivity among these mRNAs. We further demonstrate that the affinities of eIF4GI and DAP5 binding to these 5’ UTRs correlate with the efficiency with which these factors drive cap-independent translation of these mRNAs. Integrating the results of our binding and translation assays, we show that eIF4GI and/or DAP5 are critical for recruitment of a specific subset of mRNAs to the ribosome and provide mechanistic insight into their cap-independent translation.

## Introduction

Translation of mRNAs into proteins is the most energy consuming process in the cell (1,2) and plays a major role in the regulation of gene expression. In eukaryotes, initiation of cellular mRNA translation generally occurs *via* a ‘cap-dependent’ pathway in which eukaryotic initiation factor (eIF) 4E binds to the N7-methylguanosine-triphosphate (m^7^GpppN, where N is any nucleotide) ‘cap’ at the 5’ end of the mRNA to be translated (3,4). Cap-bound eIF4E subsequently recruits eIF4G, which, together with eIF4A, recruits a ribosomal 43S pre-initiation complex (PIC) composed of the 40S ribosomal subunit, a methionylated initiator tRNA (Met-tRNA_i_^Met^), and additional eIFs, to the m^7^GpppN cap. Subsequent scanning of the resulting 48S PIC to find the AUG start codon on the mRNA and joining of the 60S ribosomal subunit to the 48S PIC results in the formation of an elongation-competent 80S IC that can go on to translate the mRNA.

In addition to undergoing cap-dependent initiation, many cellular mRNAs can also initiate translation *via* ‘cap-independent’ pathways in response to changes in cellular conditions (5). The ability of these mRNAs to switch from cap-dependent to cap-independent modes of translation initiation plays an important role in maintaining normal cellular physiology (6) as well as in the cellular response to diseases such as cancer, diabetes, and, possibly, neurological disorders (7-12). A subset of cellular mRNAs, for example, has been shown to successfully bypass a global suppression of translation initiation that is caused by stress conditions such as tumor hypoxia, viral infection, nutrient deprivation, *etc*. (3,5,13) and that is driven by the sequestration of eIF4E by hypophosphorylated 4E-binding proteins (4E-BPs). While translation initiation of these mRNAs under these conditions is often referred to as cap-independent, it may be more accurately described as eIF4E-independent. Nonetheless, we will use the term cap independent here to refer to translation initiation that does not involve eIF4E-based recruitment of other eIFs to the m^7^GpppN cap. Many of the stress conditions that result in suppression of cap-dependent initiation also result in increased expression levels of eIF4GI (7,8) and/or death associated protein 5 (DAP5) (also called p97, NAT1, or eIF4G2) (14,15), suggesting that these two proteins may be involved in cap-independent initiation mechanisms.

The subset of mRNAs that is translated cap-independently under stress conditions as described above are predicted to contain highly stable structures in their 5’ untranslated regions (UTRs) that may act as internal ribosome entry sites (IRES) or cap-independent translation enhancers (CITEs) (5,16). IRES-like mechanisms involve direct recruitment of the ribosome to structured IRESs that are close to the AUG start codon (5,17), while CITE-like mechanisms involve direct recruitment of eIFs to structured CITEs near the 5’ end of the mRNA, where the eIFs are then thought to initiate cap-independent translation via 48S PIC scanning to the AUG start codon (18,19). Although chemical and enzymatic probing of cellular mRNAs thought to contain IRESs/CITEs have revealed stem loops, pseudoknots, and other structures (20,21), no common sequence or structural motifs have been identified to allow prediction of cellular IRESs/CITEs from mRNA sequence data. Regardless of whether their structured 5’ UTRs act as IRESs or CITEs, the ability of these mRNAs to bypass the 4E-BP-mediated global suppression of cap-dependent initiation has been linked to enhanced tumor development and cancer progression (2).

The overexpression of eIF4GI and DAP5 during stress conditions in which global cap-dependent translation initiation is suppressed implicates these two factors in cap-independent translation. DAP5 is a member of the eIF4G family that is homologous to the C-terminal two-thirds of eIF4GI (Fig. 1A) and that interacts with other known eIFs in manners that are both similar and distinct to that of eIF4GI. Specifically, DAP5 and eIF4GI share 39 % sequence identity in the central core region that comprises the eIF4A-, eIF3-, and RNA-binding domains (22) (Fig. 1B). Consistent with its similarity to the domain structure of eIF4GI, DAP5 interacts with eIF4A and the eIF3 component of the 43S PIC (22). Notably, however, DAP5 lacks the N-terminal, eIF4E- and polyadenine binding protein (PABP)-binding domains that are present in eIF4GI and, consequently, does not interact with eIF4E or PABP (Fig. 1A). Also, the β subunit of eIF2 (eIF2β) that interacts with the ternary complex formed by eIF2, GTP, and Met-tRNA_i_^Met^ (23) binds to the C-terminal domain of DAP5, but not to the corresponding domain in eIF4GI. Collectively, these differences between DAP5 and eIF4GI suggest differences in translation initiation mechanisms involving these two eIFs (14). For example, rather than interacting with eIF4E to recruit factors to the m^7^GpppN cap in cap-dependent initiation of mRNAs, DAP5 has instead been shown to mediate the cap-independent initiation of a subset of mRNAs, including those encoding Bcl2, Apaf-1, p53, and DAP5 itself (24,25). With the exception of p53 mRNA, which contains a structural element that functions as an IRES and has been shown to bind DAP5 using electrophoretic mobility shift assay (EMSA) studies (24), there have not yet been any studies aimed at investigating whether mRNAs translated cap-independently directly recruit eIF4GI and/or DAP5 to the 5’ UTRs and how such recruitments might drive the switch from cap-dependent to cap-independent initiation in response to cellular stress.

**Figure 1.**
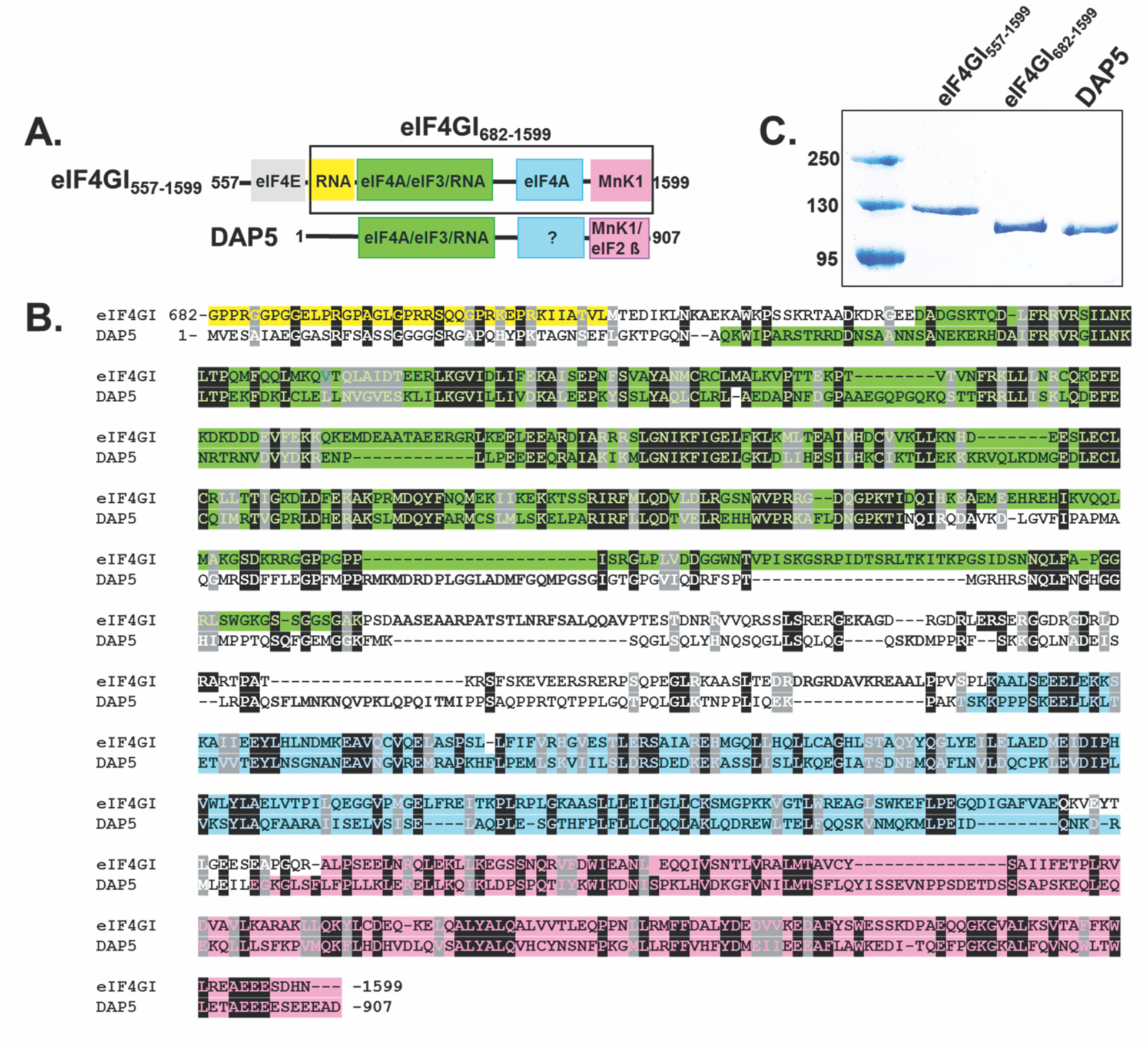
Domain structure and sequence alignment between eIF4GI and DAP5 **A**. Cartoons showing the domain architecture of eIF4GI_557-1599_ and full length DAP5. The shorter construct of eIF4GI, eIF4GI_682-1599,_ is highlighted in the black box. **B**. Sequence alignment encompassing similar domains in eIF4GI_682-1599_ and DAP5. The sequence alignment was performed using T-coffee (59). Residues are color-coded according to their domain organization shown in **A. C**. A 10 % SDS-PAGE gel showing purity of eIF4GI_557-1599_, eIF4GI_682-1599_, and DAP5 used for this study.

To address these gaps in our understanding, we used a fluorescence anisotropy-based equilibrium binding assay to measure the affinities with which two variants of human eIF4GI, one that lacks the N-terminal, eIF4E-binding domain (eIF4GI_682-1599_) and one that contains the eIF4E binding domain (eIF4GI_557-1599_), as well as full-length human DAP5, bind to RNA oligonucleotides corresponding to the 5’ UTRs of a representative set of mRNAs. These mRNAs bypass (2) the 4E-BP-mediated global suppression of cap-dependent initiation and encode HIF-1α, FGF-9, and p53. These mRNAs were chosen because HIF-1α and FGF-9 have been shown to be translationally upregulated in hypoxic conditions where cap-dependent translation is suppressed and where eIF4GI and 4EBPI levels are overexpressed (2,26). p53 mRNAs were chosen because these mRNAs are known to interact with DAP5 as part of a cap-independent translation mechanism (24). Complementing these binding assays, a luciferase-based gene expression reporter assay was used to characterize whether and to what extent binding of eIF4GI_682-1599_, eIF4GI_557-1599_, and/or DAP5 promotes the translation of luciferase-encoding mRNAs containing these same 5’ UTRs (UTR-Luc mRNAs) in a rabbit reticulocyte lysate-based cap-dependent *in vitro* translation system. Using this assay with UTR-Luc mRNA constructs that either lack or have a highly stable, engineered stem-loop near the 5’ end of the 5’ UTR that blocks initiation from a free 5’ end, has further allowed us to investigate whether these mRNAs use an IRES-like or CITE-like mechanism of translation initiation. The results of our experiments demonstrate that eIF4GI_682-1599_, eIF4GI_557-1599_, and DAP5 exhibit differential binding affinities to the 5’ UTRs of the mRNAs encoding HIF-1α, FGF-9, and p53; that the eIF4E-binding domain of eIF4GI confers additional binding affinity and specificity; and that binding affinity positively correlates with the abilities of these eIFs to drive translation of these same UTR-Luc mRNAs. Moreover, the inhibition, or lack thereof, of translation initiation by the engineered stem-loop suggests that some of these mRNAs use an IRES-like mechanism of translation initiation which does not require a free 5’ end, while others use a CITE-like mechanism (27). Based on our collective results, we show mRNAs with structured 5’ UTRs that act as IRESs or CITEs selectively recruit eIFs for cap-independent initiation and drive expression of the proteins they encode.

## Results

### eIF4GI and DAP5 bind specifically and with differential affinities to the 5’ UTRs of a subset of cellular mRNAs

To characterize the binding of eIF4GI and DAP5 to the 5’ UTRs of the mRNAs encoding HIF-1α, FGF-9, and p53, we used a fluorescence anisotropy-based equilibrium binding assay developed in our laboratories (Fig. 2A). This assay produces binding curves from fluorescence anisotropy changes that arise as titrated proteins bind to RNA oligonucleotides that are covalently, 5′-end labeled with fluorescein and that lack a m^7^GpppN cap. Four RNA oligonucleotides comprising the 5’ UTRs of mRNAs encoding HIF-1α, FGF-9, and the two 5’ UTRs of p53, were assayed. The two p53 5’ UTRs represent the two 5’ UTRs involved in translation of the two distinct isoforms of p53. Each p53 5’ UTR contains elements that select for translation initiation at either an upstream start codon (p53_A_) or a downstream start codon (p53_B_), which produce the full-length p53 (FL-p53) or an N-terminal-truncated isoform (ΔN-p53), respectively (28). When eIF4GI_682-1599_, eIF4GI_557-1599_, or DAP5 binds to one of the fluorescein-labeled 5′ UTRs, there is an increase in molecular weight of the 5’ UTR that increases the rotational time of the 5’ UTR. The slower rotation is observed as increased fluorescence anisotropy of the fluorescein reporter. To quantify the affinity of eIF4GI_557-1599_, eIF4GI_557-1599_, and DAP5 binding to the four 5′ UTRs, we recorded the change in the fluorescence anisotropy of each fluorescein-labeled 5’ UTR as a function of increasing concentrations of eIF4GI_682-1599_, eIF4GI_557-1599_, and DAP5 and fitted the resulting data points with a single-site, equilibrium-binding isotherm (Fig. 2B, 2C and 2D). The results of these experiments (Table 1) demonstrate that eIF4GI_682-1599_, eIF4GI_557-1599_, and DAP5 bind to the four 5′ UTRs with equilibrium dissociation constants (*K*_d_s) ranging from 12–290 nM (Table 1).

**Table 1.** Parameters describing the equilibrium binding of eIF4GI constructs and DAP5 to the 5’ UTRs.

| 5' UTRs | eIF4GI<sub>557-1599</sub> |  |  | eIF4GI<sub>682-1599</sub> |  |  | DAP5 |  |  |
| --- | --- | --- | --- | --- | --- | --- | --- | --- | --- |
| | $K_d \pm \text{S.D.}$<br>(nM)<sup>a</sup> | Amp.<br>($r_{\text{max}} - r_{\text{min}}$)<sup>b</sup> | $\chi^2$<sup>(c)</sup> | $K_d \pm \text{S.D.}$<br>(nM) | Amp.<br>($r_{\text{max}} - r_{\text{min}}$) | $\chi^2$ | $K_d \pm \text{S.D.}$<br>(nM) | Amp.<br>($r_{\text{max}} - r_{\text{min}}$) | $\chi^2$ |
| HIF1α | 50 ± 6.0 | 0.045 | 0.996 | 139 ± 23.0 | 0.034 | 0.994 | 154 ± 18.0 | 0.061 | 0.988 |
| FGF-9 | 12 ± 2.0 | 0.079 | 0.995 | 120 ± 7.0 | 0.027 | 0.996 | 136 ± 10.0 | 0.039 | 0.998 |
| p53<sub>A</sub> | 86 ± 4.0 | 0.021 | 0.998 | 248 ± 16.0 | 0.036 | 0.999 | 290 ± 14.0 | 0.044 | 0.999 |
| p53<sub>B</sub> | 68 ± 19.0 | 0.025 | 0.998 | 126 ± 4.0 | 0.017 | 0.998 | 112 ± 6.0 | 0.048 | 0.998 |

<sup>a</sup> $K_d$ is the equilibrium dissociation constant<sup>b</sup> $r_{\text{max}} - r_{\text{min}}$ is the amplitude which indicates change in anisotropy<sup>c</sup> $\chi^2$ is the goodness of fit

**Table 2.**
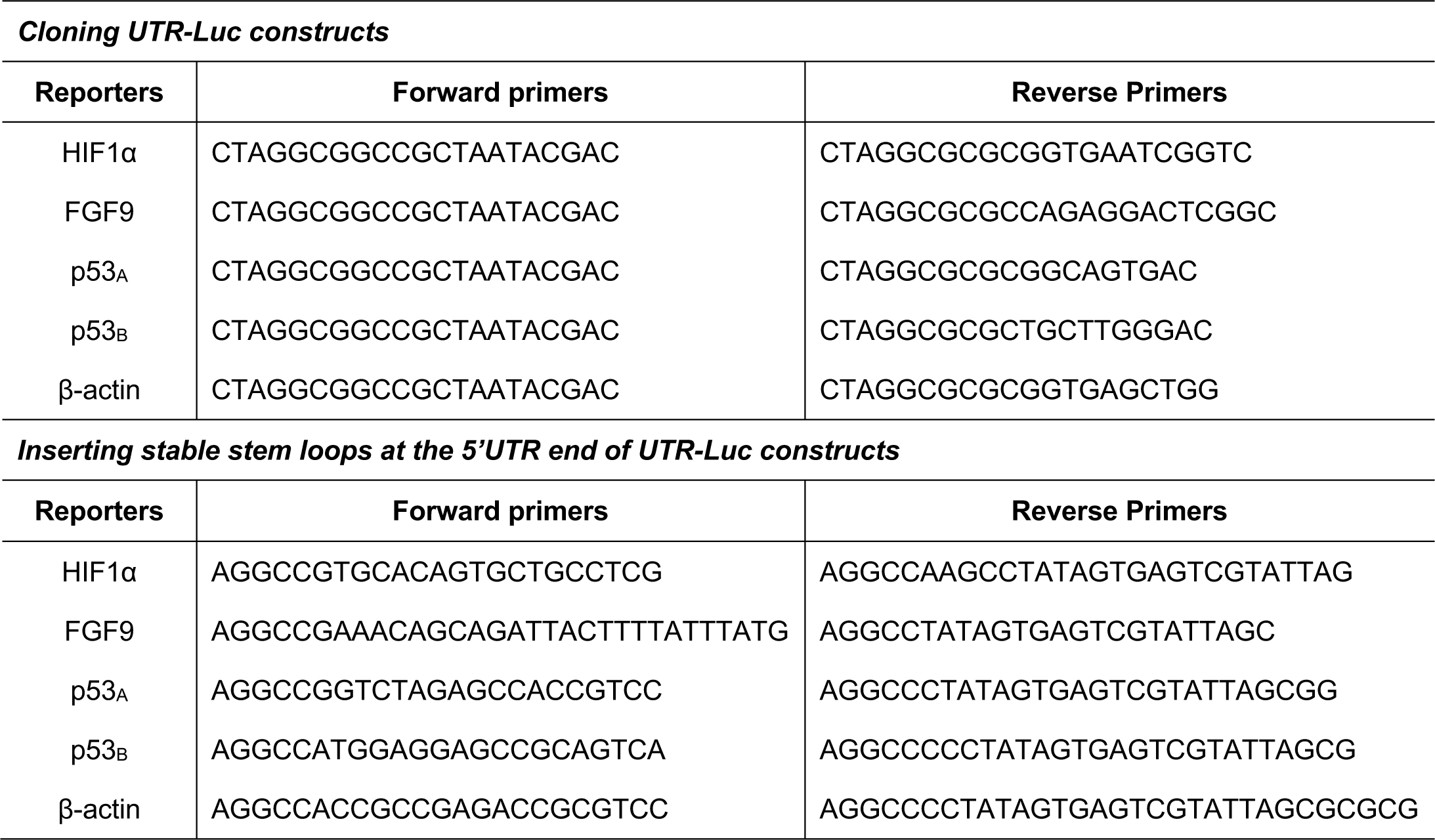
Primers used for cloning UTR-Luc constructs and inserting stable stem-loops at the 5’ UTR of UTR-Luc constructs

**Figure 2.**
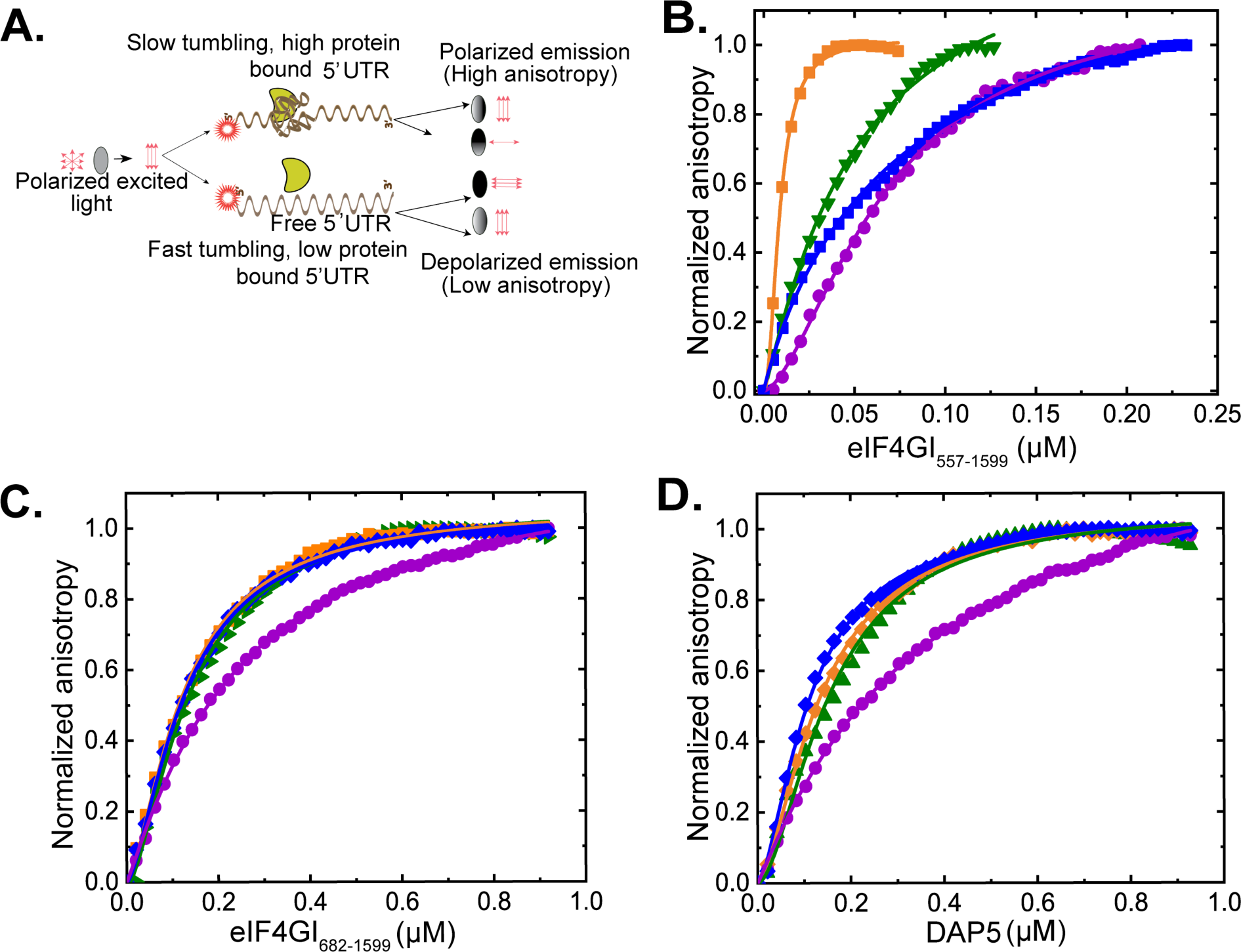
Equilibrium binding-titrations of 5’ UTRs with eIF4GI_557-1599_, eIF4GI_682-1599_, and DAP5. **A**. Cartoon showing the fluorescence anisotropy-based equilibrium binding assay. Normalized anisotropy changes for the interaction of fluorescein-labeled FGF-9 (-▪-), HIF-1α (-▾-), p53_A_ (-•-) and p53_B_ (-♦-) 5’ UTRs, with **B**. eIF4GI_557-1599_, **C**. eIF4GI_682-1599_, and **D**. DAP5. 100 nM of fluorescein-labeled 5’ UTR in Titration Buffer was titrated with increasing concentrations of eIF4GI_557-1599_, eIF4GI_682-1599_, or DAP5 at 25 °C and the anisotropy at each titration point was measured using excitation and emission wavelengths of 495 nm and 520 nm, respectively. Data points correspond to the average of three independent anisotropy measurements and the curves represent the non-linear fits that were used to obtain the averages and standard deviations for the corresponding K_d_ values.

Comparative analyses of our results demonstrate that eIF4GI_682-1599_, eIF4GI_557-1599_, and DAP5 exhibit differential binding affinities among the 5′ UTRs. Specifically, eIF4GI_682-1599_ binds to the p53_A_ 5′ UTR with a *K*_d_ that is ∼2-fold higher (i.e., binding that is ∼2-fold weaker) than that with which it binds to the other 5’ UTRs (Table 1). DAP5 exhibited an even greater difference in binding affinity among the 5′ UTRs, with a *K*_d_ for the p53_B_ 5′ UTR that is more than 2.5-fold lower than the *K*_d_ for the p53_A_ 5′ UTR. It is important to note, however, that the trend of the differences in binding affinities between the two translation factors (eIF4GI_682-1599_ and DAP5) for the 5′ UTRs is similar. However, dramatic differences in binding among the 5’ UTRs were observed for eIF4GI_557-1599_. Inclusion of the eIF4E-binding domain in eIF4GI_557-1599_ relative to eIF4GI_682-1599_ increased the binding affinity of eIF4GI_557-1599_ from 1.8-fold (p53_B_ 5’ UTR) to 10-fold (FGF-9 5’ UTR) compared to those measured for eIF4GI_682-1599_. Further, the differences in binding affinity among the 5’ UTRs were as much as 7-fold different compared to an approximately 2-fold difference among the 5’ UTRs for eIF4GI_682-1599_. Taken together, these results suggest much of the specificity of eIF4GI binding to the 5’ UTRs is conferred by the eIF4E-binding domain (i.e., residues 557-682) of eIF4GI.

### The binding of eIF4GI and DAP5 is specific to the 5’ UTRs of the selected mRNAs

In order to further investigate whether binding of eIF4GI and DAP5 to RNA depends on structural features within the RNA and to contextualize and validate the results of the binding studies above, we assessed the binding of eIF4GI_682-1599_, eIF4GI_557-1599_, and DAP5 to a presumably unstructured, 101-nucleotide poly(UC) RNA oligonucleotide and an oligonucleotide encompassing the 5’ UTR of the mRNA encoding β-actin, which has been reported to utilize a cap-dependent mechanism for translation initiation (29). The results of these experiments demonstrate that eIF4GI_682-1599_, eIF4GI_557-1599_, and DAP5 do not exhibit appreciable binding to the polyUC oligonucleotide (Fig. 3A, B and C and Table S1) nor to the β-actin 5’ UTR, findings consistent with the hypothesis that structural features within the 5’ UTRs of our subset of RNAs act as specific recognition elements and binding sites for eIF4GI and DAP5.

**Figure 3.**
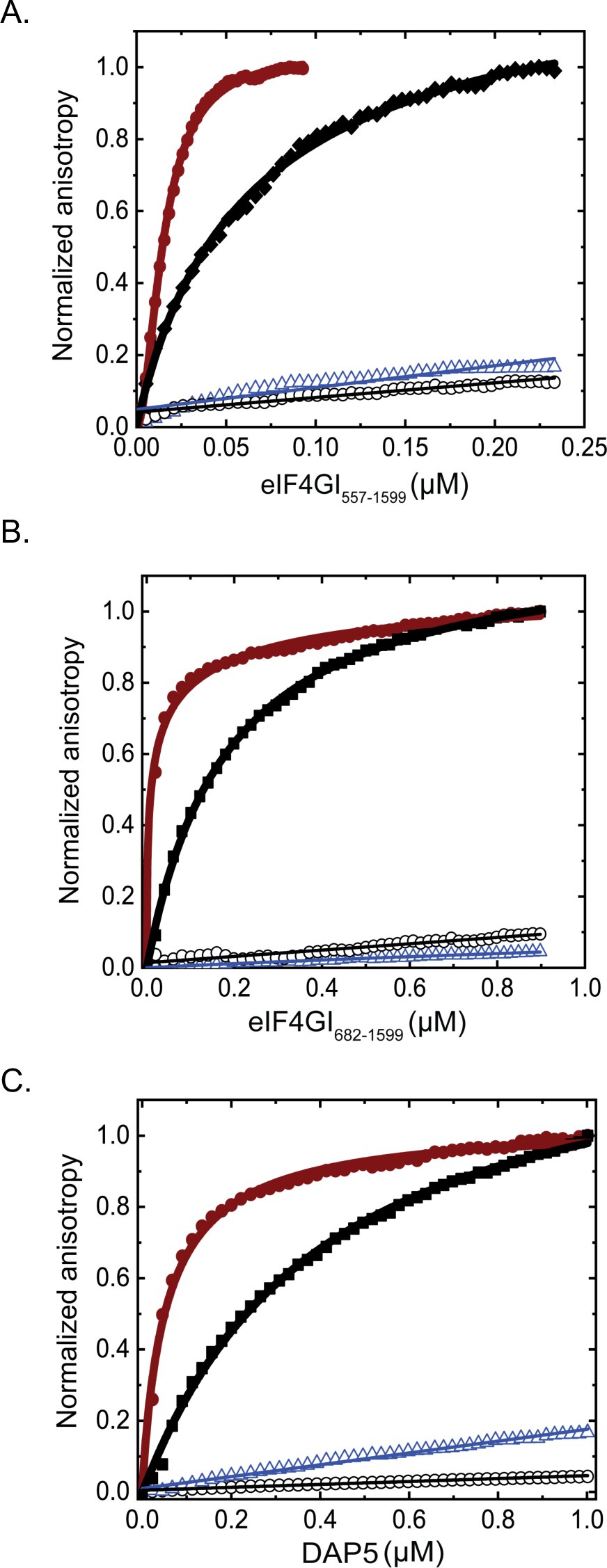
Equilibrium binding titrations of fluorescein-labeled ferritin IRE (-•-), EMCV J/K IRES (-▪-), β-actin UTR (-°-), and polyUC (-▽-) with **A**. eIF4GI_557-1599_, **B**. eIF4GI_682-1599_, and **C**. DAP5. Average of 3 independent experiments were performed and data were collected and analyzed as Fig. 2.

Previously, we have shown that eIF4F (a complex composed of eIF4GI, eIF4A, and eIF4E) binds to the 30-nucleotide iron responsive element (IRE) stem-loop within the 5’ UTR of the mRNA encoding ferritin with a *K*_d_ of 9 nM, a binding interaction that stimulates ferritin mRNA translation in response to elevated cellular concentration of iron (30). Using the fluorescence anisotropy-based equilibrium binding assay, here we show that eIF4GI_682-1599_, eIF4GI_557-1599_, and DAP5 bind to the IRE with *K*_d_s of 18 nM, 15 nM and 35 nM, respectively (Fig. 3 and Table S1). Similarly, we performed experiments in which we quantified the affinity of eIF4GI_682-1599_, eIF4GI_557-1599_, and DAP5 for the J/K domain of the IRES in the 5’ UTR of the positive strand genomic RNA, encoding the encephalomyocarditis virus (EMCV) polyprotein (Fig. 3 and Table S1). These experiments show that eIF4GI_682-1599_, eIF4GI_557-1599_, and DAP5 bind to the EMCV J/K IRES with *K*_d_s of 175 nM, 63 nM, and 519 nM, respectively. The *K*_d_ for binding of eIF4GI_682-1599_ to EMCV J/K IRES RNA is similar to what has been previously reported for the binding of human eIF4GI (643-1076) to the EMCV J/K IRES using EMSA (170 nM, (31)) as part of the mechanism through which eIF4GI drives expression of the EMCV polyprotein in human cells (32). The same report showed that a construct containing an N-terminal portion of DAP5 (62-330) did not bind to the EMCV J/K IRES, consistent with our much higher K_d_ (519 nM) for full-length DAP5 compared to eIF4GI constructs binding.

### The 5’ UTRs of a subset of cellular mRNAs that bind eIF4GI and DAP5 can drive cap-independent translation

Having demonstrated that eIF4GI and DAP5 bind specifically and with relatively high affinity to the 5’ UTRs of our subset of mRNAs, we examined the cap-independent translation of a set of reporters containing these 5’ UTRs. To quantify the cap-independent activities of our selected mRNAs, the 5’ UTRs of the HIF-1α, FGF-9, p53_A_, and p53_B_ mRNAs and, as a control, the 5’ UTR of β-actin mRNA were cloned upstream of a luciferase reporter gene (Fig. 4A). The resulting capped, polyadenylated UTR-Luc mRNAs were translated using a nuclease-treated, rabbit reticulocyte (RRL)-based, cap-dependent, *in vitro* translation system containing a natural abundance of the eIFs. To assess cap-independent luciferase expression, a non-functional cap analog (ApppG) was used to cap the UTR-Luc mRNAs (hereafter referred to as the ApppG-capped transcripts). The cap-independent expression levels of the ApppG-capped transcripts were compared with the expression observed when these transcripts were capped with a functional m^7^GpppA cap. We observed that the activities of the ApppG-capped transcripts ranged from ∼70 % for FGF-9 UTR-Luc mRNA to 8 % for p53_B_ UTR-Luc mRNA when compared to their corresponding m^7^GpppA-capped transcripts (Fig. S1). Expression of the ApppG-capped β-act-UTR-Luc mRNA was reduced to less than 2 % of the corresponding m^7^GpppA-capped construct (Fig. S1). These results further support the notion that the 5’ UTR of a subset of cellular mRNAs can drive cap-independent translation, albeit at a relatively lower efficiency than cap-dependent translation. During stressful conditions in which cap-dependent translation is inhibited and expression of eIF4GI and DAP5 are elevated, expression from these 5’ UTRs can help cells mitigate the effects of the stressor (2).

**Figure 4.**
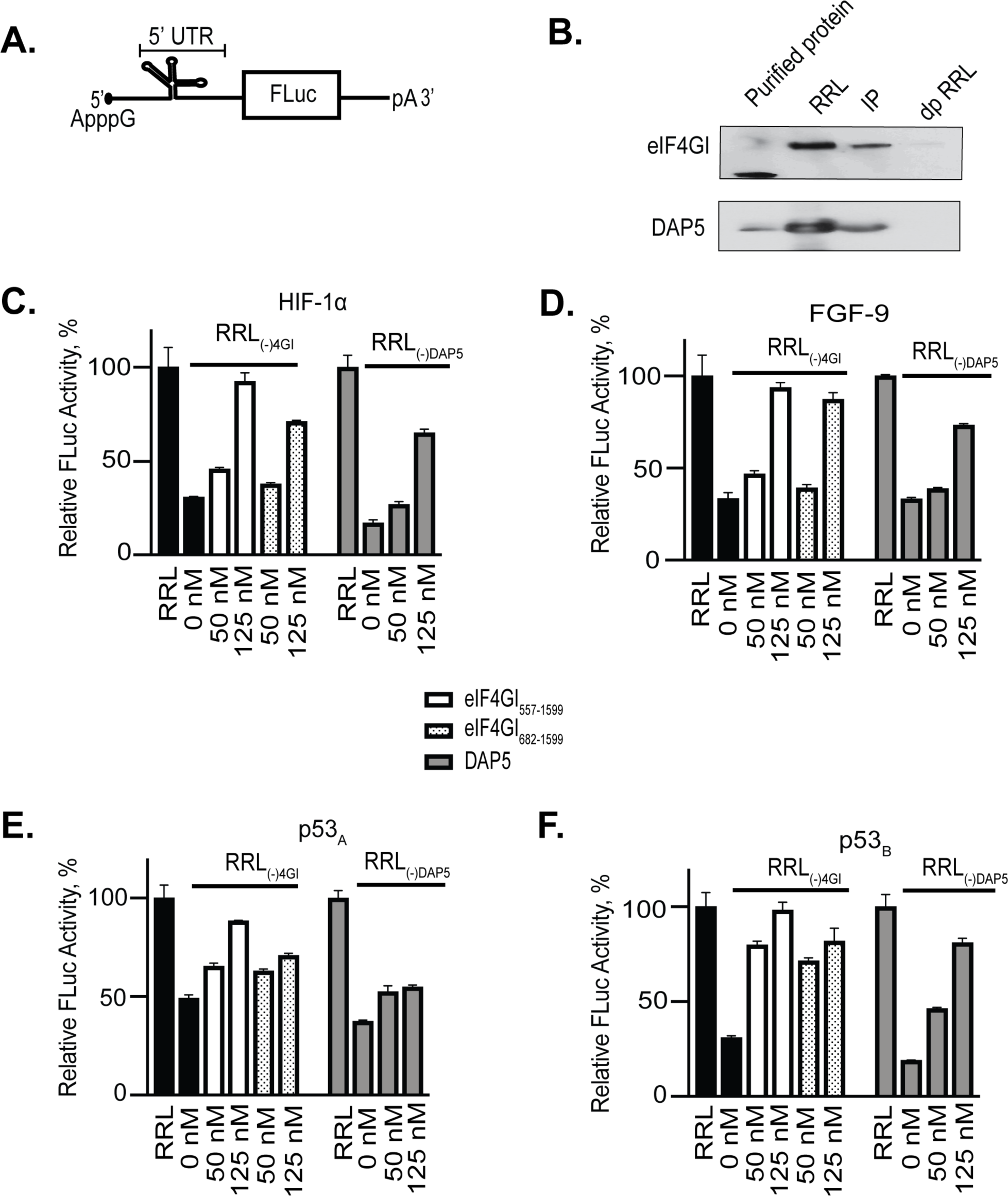
Effect of eIF4GI_557-1599_, eIF4GI_682-1599_, and DAP5 on the expression of UTR-Luc mRNAs. **A**. Cartoon showing the design of UTR-Luc mRNA reporters construct containing a structural element upstream of the firefly luciferase gene. **B**. Western blot of eIF4GI and DAP5 depleted RRL probed using specific eIF4GI and DAP5 antibodies. Purified eIF4GI_682-1599_ or DAP5 protein used for the study were included as positive controls (Lane 1). Lane 2 shows non-depleted RRL; Lane 3 immuno-precipitate pulled down by antibodies for either eIF4GI (top panel) or DAP5 (lower panel). dpRRL is the depleted RRL (RRL_(–)4GI_ or RRL_(–)DAP5_). Lysates were run on 4-12 % SDS gradient gels and transferred to a nitrocellulose membrane. The effect of increasing concentrations of eIF4GI_557-1599_, eIF4GI_682-1599_, and DAP5 on the translation yields of ApppG-capped transcripts of **C**. HIF-1α, **D**. FGF-9, **E**. p53_A_, and **F**. p53_B_ UTR-Luc-mRNAs. Relative luciferase activity measured in RRL (Bar 1 or Bar 7) or RRL_(–)4GI_ or RRL_(–)DAP5_ (Bar 2 or Bar 8) in the presence of increasing concentrations of either eIF4GI_557-1599_ (Bar 3-4), eIF4GI_682-1599_ (Bar 5-6), and DAP5 (Bar 9-10). Relative luciferase activity was normalized to the control reaction (ApppG-capped UTR-Luc mRNA) for each reporter performed in RRL. Bar heights and error bars correspond to the average and standard deviations, respectively, of three relative luciferase activity measurements and t-tests using the average and standard deviation of all experiments exhibited a p < 0.001.

In order to further support the idea that translation initiation of these mRNAs was independent of eIF4E, we determined the extent to which translation of the m^7^GpppA- and ApppG-capped transcripts were affected by 4EGI-1, a small-molecule that binds to the same site on eIF4E that interacts with eIF4GI and inhibits the interaction of eIF4GI and eIF4E (33). Cap-dependent initiation was significantly suppressed through the addition of 4EGI-1 to m^7^GpppA-capped β-act-UTR-Luc mRNA, but had no effect on the ApppG-capped transcripts of our selected 5’ UTRs, with the exception of the ApppG-capped HIF-1α transcript, which exhibited an ∼44 % reduction in translation (Fig. S2). These results show that, to a large extent, translation of the ApppG-capped transcripts proceeds *via* an eIF4E-independent mechanism. The observation that the ApppG-capped HIF-1α transcript expression is partially reduced suggests that eIF4E might not function exclusively during cap-dependent translation initiation, but may also play an important role in the cap-independent translation initiation of some mRNAs. For example, eIF4E has been shown to stimulate the helicase activity of eIF4A in the case of some highly structured 5’ UTRs (34), such as HIF-1α, and/or to induce conformational changes in eIF4G that enhance its binding to mRNAs (35).

### eIF4GI and DAP5 stimulate and restore cap-independent in vitro translation of the selected 5’ UTR-Luc mRNAs

Having established that eIF4GI and DAP5 bind specifically and with relatively high affinity to the HIF-1α, FGF-9, p53_A_, and p53_B_ 5’ UTRs and that the corresponding UTR-Luc mRNAs are translated through a cap-independent pathway, we sought to establish the extent to which this cap-independent translation initiation of these mRNAs was dependent on eIF4GI and DAP5. To accomplish this, we used our luciferase-based reporter assay and first determined the effects on translation of depletion of either eIF4GI or DAP5. We measured the luciferase activity produced by each ApppG-capped transcript in an RRL that had been depleted of either endogenous eIF4GI (RRL_(–)4GI_) or DAP5 (RRL_(–)DAP5_) using antibodies directed against these proteins. Western blots confirmed successful depletion of eIF4GI or DAP5 (Fig. 4B) from the RRLs. As a control to test for the indirect depletion of other eIFs, eIF4E, eIF2β, and eIF4A levels were tested in RRL_(–)4GI_ or RRL_(–)DAP5_ and were found to be similar to, or only slightly reduced from, those seen in RRL (Fig. S3). Using either RRL_(–)4GI_ or RRL_(–)DAP5_, we measured the luciferase activity produced by each translation reaction of ApppG-capped transcript and compared it to RRL translation assays of the same transcript without depletion of endogenous eIF4GI or DAP5. To confirm the eIF4GI or DAP5 dependence on the translation initiation of these UTR-Luc mRNAs, we added increasing concentrations of exogenous eIF4GI_557-1599_, eIF4GI_682-1599_, or DAP5 to the corresponding depleted RRL. Translation of UTR-Luc mRNAs were significantly reduced in the depleted RRLs compared with the RRL control (Fig. 4 C-F). Specifically, the translation output of HIF-1α, FGF-9, p53_A_, and p53_B_ ApppG-capped UTR-Luc mRNAs were reduced by 70 %, 67 %, 51 %, and 70 %, respectively, in RRL_(–)4GI_ (Fig. 4C-F). Addition of eIF4GI_557-1599_ or eIF4GI_682-1599_, significantly rescued and stimulated the cap-independent translation initiation of all four UTR-Luc mRNAs (Fig. 4 C-F). For RRL_(-)4GI_, eIF4GI_557-1599_ restored translation of the ApppG-capped transcripts to ∼85-100 % of the levels observed in RRL whereas eIF4GI_682-1599_ was slightly less effective, restoring levels to ∼70-90 % of the RRL levels for the same transcripts (Fig 4 C-F). In the case of RRL_(-)DAP5_, the translation outputs of HIF-1α, FGF-9, p53_A_, and p53_B_ ApppG-capped UTR-Luc mRNAs were reduced by 83 %, 67 %, 63 %, and 81 %, respectively. Addition of DAP5 was also able to rescue cap-independent translation initiation of all four UTR-Luc mRNAs, although with somewhat less efficiency compared to the eIF4GI constructs. DAP5 restored translation to ∼45-65 % of the levels for the same transcripts in RRL (Fig. C-F). These data follow the same overall trend as our fluorescence anisotropy-based binding assay data where eIF4GI_557-1599_ showed the highest binding affinity and DAP5 the lowest affinity to the 5’ UTRs (Table 1).

Further, the ApppG-capped β-act-UTR-Luc mRNA showed less than 2 % of the translation output as compared to the corresponding m^7^GpppA-capped transcript. In agreement with our binding results, eIF4GI_682-1599,_ eIF4GI_557-1599_, and DAP5 did not show any significant stimulation of the cap-independent translation of this ApppG-capped β-act-UTR-Luc mRNA, results that are consistent with the previous and widespread use of this mRNA as a control for cap-dependent initiation and translation (29).

### A free 5’ end is important for the cap-independent translation activities of HIF-1α and p53_A_ UTR-Luc mRNAs, but not FGF-9 and p53_B_ UTR-Luc mRNAs

In order to gain a better mechanistic understanding of the cap-independent translation initiation of our selected mRNAs, we introduced a highly stable stem-loop at the 5’ end of our mRNAs (Fig. 5A). This same stem-loop at this same position has been previously used to block scanning from the free 5’ end of an mRNA while having no effect on the internal translation initiation mediated by an IRES (25,27,36). We found that the stem-loop structure significantly repressed translation of m^7^GpppA-capped β-act-UTR-Luc mRNA (Fig. 5B), which served as a positive control for inhibition of 5’-end-scanning-dependent translation initiation. Comparative analyses of our selected UTR-Luc mRNAs showed that the stem-loop did not affect the translation initiation activities of FGF-9 UTR***-***Luc mRNA (Fig. 5D) and p53_B_ UTR***-***Luc mRNA (Fig. 5F), suggesting that internal initiation occurred on these UTR-Luc mRNAs. In contrast, the translation efficiencies of HIF-1α Luc mRNA (Fig. 5C) and p53_A_ UTR-Luc mRNA (Fig. 5E) were significantly reduced by the stem-loop, suggesting that translation of these UTR-Luc mRNAs required a free 5’ end for translation initiation and that these 5’ UTRs contain structural elements that may act as CITEs rather than IRESs.

**Figure 5.**
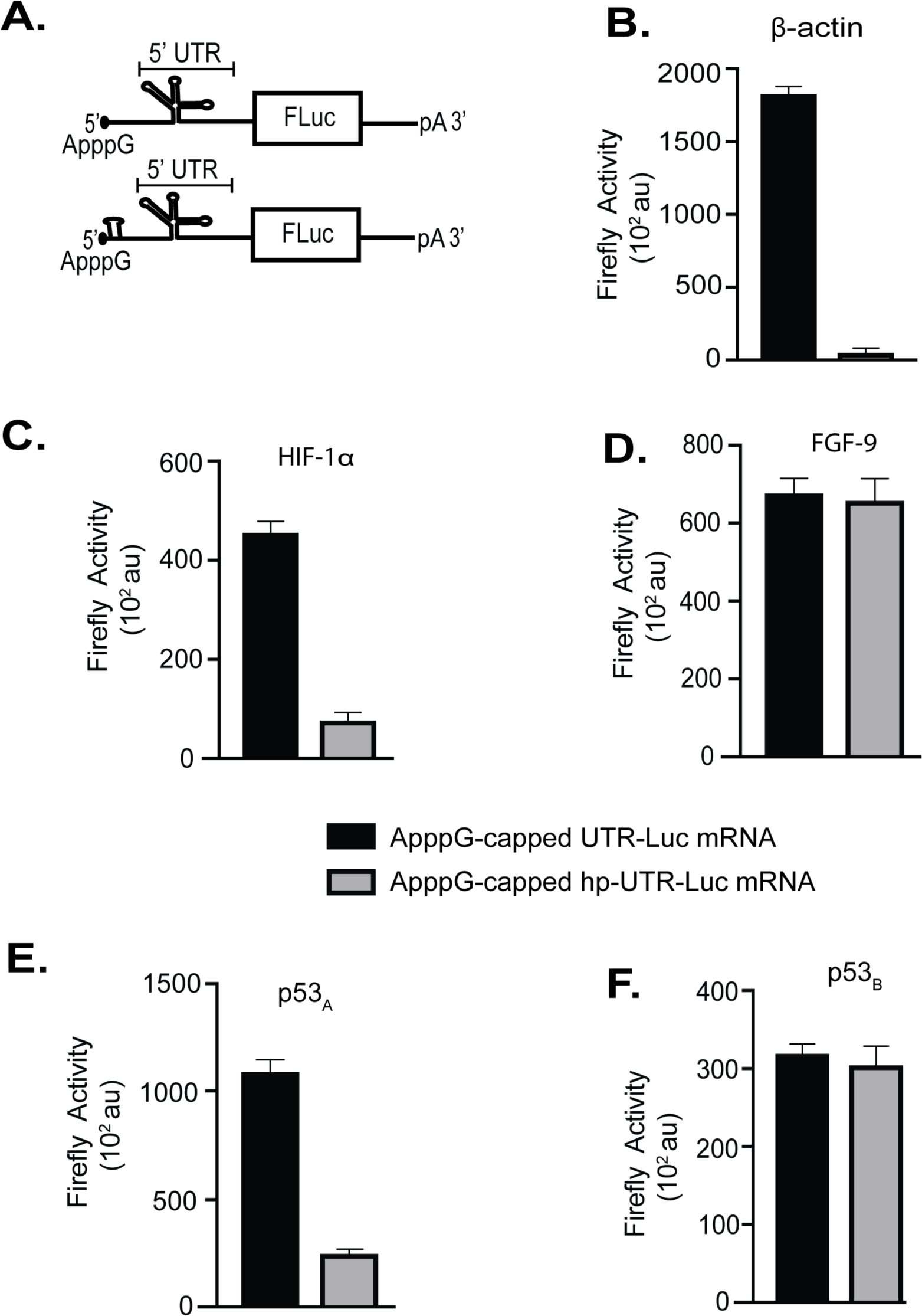
Effect of a stable 5’ hairpin loop inserted at the 5’ terminal of UTR-Luc mRNAs. **A**. Cartoon representation of the reporter construct used to identify IRES or CITE elements within the 5’ UTRs of HIF-1α, FGF-9, p53_A_, and p53_B_ UTR-Luc mRNAs. **B**. Comparison of the translation output of reporters ApppG-capped UTR-Luc mRNAs *versus* ApppG-capped hp-UTR-Luc mRNA.

## Discussion

In this study, using a fluorescence anisotropy-based equilibrium binding assay and a luciferase-based gene expression reporter assay, we have demonstrated that eIF4GI and DAP5 directly bind to and stimulate cap-independent translation initiation of a subset of cellular mRNAs. The truncated eIF4GI proteins, eIF4GI_557-1599_ and eIF4GI_682-1599_, as well as DAP5 bind to the 5’ UTRs of this subset of mRNAs with *K*_d_s in the range of 12-290 nM (Fig. 2, 3 and Table 1) and stimulate cap-independent translation of the corresponding UTR-Luc mRNAs by 2-5 fold over lysates depleted of either eIF4GI or DAP5 (Fig. 4 C-F). These *K*_d_s are comparable to those observed for the binding of eIF4GI constructs to the EMCV J/K IRES (Fig. 3 and Table S1), a well characterized system in which it has been previously shown that binding of truncated eIF4GI_643-1076_ to a well-defined structural feature within the 5’ UTR of EMCV mRNA stimulates the cap-independent translation of this mRNA (31). In line with cellular data (2,15), our observations strongly suggest that both eIF4GI and DAP5 recognize and bind to specific structural features within the 5’ UTRs of our selected mRNAs. Moreover, the observation that eIF4GI_682-1599_ and DAP5 bound to our selected 5’ UTRs with similar binding affinities (Fig. 2 and Table 1) suggest that eIF4GI and DAP5 recognize similar elements within the various 5’ UTRs. Furthermore, the intriguing differences between the *K*_d_s for the larger construct of eIF4GI_557-1599_ and DAP5 (Table 1) suggest that, to bring about translational outcomes, cellular mRNAs with structured 5’ UTRs may preferably recruit eIF4GI, depending on its cellular availability, which may change by the type of stress conditions, cell-type, or stability of the proteins (2,37,38).

Several reports have demonstrated that a subset of cellular mRNAs possess IRES-like elements in addition to the cap structure, and that these mRNAs act to recruit ribosomal PICs and presumably key eIFs internally to the 5’ UTR of these mRNAs in order to initiate cap-independent translation (2,5,39). However, no common structural motifs were identified among the cellular IRES elements and, compared to viral IRES, cellular IRES elements appear to be much more diverse and less stable in terms of Gibbs free energy of folding (40). In contrast to these reports, it has recently been proposed that other cellular mRNAs may recruit ribosomal PICs in an cap-independent manner to the 5’ end using a CITE that recruits eIFs, and that the ribosomal PIC then scans from the 5’ end rather than using an IRES to initiate translation internally (19). Given the fact that some 5’ UTRs may possess either IRESs and/or CITEs, we were prompted to test the hypothesis that the free 5’ end of our selected 5’ UTRs is required for translation. Our data showed that the presence of a highly stable, scanning-inhibiting stem-loop (36,41) at the 5’ end of the FGF-9 and p53_B_ ApppG-capped hp-UTR-Luc mRNAs did not affect the absolute translation levels of these mRNAs (Fig. 5), demonstrating that they were cap-independently translated, presumably using IRES-like mechanisms. In contrast with this, the stem-loop at the 5’ end of the HIF-1α and p53_A_ ApppG-capped hp-UTR-Luc mRNAs repressed the cap-independent translation initiation activities of these UTR-Luc mRNAs (Fig. 5), suggesting that accessibility of the 5’ end of these cellular mRNAs is necessary and, correspondingly, a CITE-like mechanism is important for the initiation of these mRNAs. Because the subset of cellular mRNAs we have investigated here employ different systems of cap-independent translation, we speculate that this phenomenon may provide additional regulation for the expression of these mRNAs.

Our findings are reminiscent of the manner in which viruses use highly structured IRESs to directly recruit eIFs, the 40S subunit, and/or the 43S PIC to drive the cap-independent translation of viral mRNA (5,42,43). Although the mechanisms through which viral mRNAs use IRESs to drive cap-independent initiation have been extensively characterized using genetic, biochemical, and structural approaches (16,44,45), the mechanisms through which eukaryotic cellular mRNAs drive cap-independent initiation remain largely unknown. Nonetheless, an increasing body of evidence suggests eukaryotic cellular mRNAs can employ non-canonical initiation mechanisms that are distinct from those employed by IRES-containing viral mRNAs. For example, Cate and co-workers have recently shown that the ‘d’ subunit of eIF3 (eIF3d) targets mRNAs encoding proteins involved in cell proliferation and serves as a transcript-specific, cap-binding protein (46). Using the mRNA encoding the *c-Jun* transcription factor as an example, these authors showed that, under conditions in which an RNA structure in the 5’ UTR blocks eIF4E from binding to the 5’ cap, eIF3d binds directly to the 5’ cap and serves as an alternative cap-binding factor (47). Even more recently, it has been shown that DAP5 is a direct binding partner of eIF3d and it has been proposed that DAP5 uses eIF3 to direct the eIF4E-independent, cap-dependent translation of a subset of cellular mRNAs when cellular stress conditions lead to inactivation of eIF4E (48). In addition to these mechanisms in which RNAs utilize IRES elements, eIF3d, or other eIFs that function as alternative cap-binding proteins, the methylation of adenosine residues in the 3’- and 5’ UTRs of eukaryotic cellular mRNAs have been shown to stimulate translation by an unknown mechanism (49,50). The cellular IRES- and CITE-based mechanisms we have investigated here are distinct from these other types of mechanisms that make use of an alternative cap-binding protein or post-transcriptional modification of the mRNA to be translated. Here, eIF4GI or DAP5 completely bypass any cap-dependent processes and are instead directly recruited to IRESs or CITEs within the mRNAs. Indeed, because the binding studies we present here employ purified eIF4GI_557-1599_, eIF4GI_682-1599_, and DAP5 and uncapped RNA oligonucleotides, our results show these translation factors can bind directly to 5’ UTRs of select mRNAs in a completely cap-independent manner and without the need of alternative cap-binding proteins or mRNA methylation. Similarly, because the gene expression assays we present here also employed purified mRNAs with non-functional cap analogs and are performed in RRL, RRL_(-)4GI_ and RRL_(-)DAP5_ we can be confident that the stimulation of translation we observe is due to the direct interaction of eIF4GI or DAP5 with the IRESs or CITEs in the 5’ UTR of our selected mRNAs, rather than to the indirect effects of alternative cap-binding proteins or methylation of the mRNA.

Based on our observations, we propose that an important initial event in cap-independent initiation of eukaryotic cellular IRES- or CITE-containing mRNAs is binding of eIF4GI or DAP5, similar to the cap-independent initiation of CITE- or IRES-containing viral mRNAs (51-53). Specifically, we propose that eIF4GI or DAP5, perhaps with the aid of additional factors, bind to elements within the 5’ UTRs and subsequently recruit additional eIFs, the 40S subunit, and/or the 43S PIC to these mRNAs. In analogy to the cap-independent initiation of several CITE-containing viral mRNAs (51,53,54) we propose that the resulting 48S PIC may then scan to the AUG start codon. Our data suggest that this is the case for the for HIF-1α and p53_A_ mRNA. Alternatively, these factors may be bound to elements near the AUG acting as IRES-like structures as appears to be the case for FGF-9 and p53_B_ (Fig. 6). Given the current inability to predict IRESs or CITE elements from sequence data for cellular mRNAs, it is not surprising that multiple mechanisms and eIF requirements may be present. This is certainly the case for viral cap-independent translation (43).

**Figure 6.**
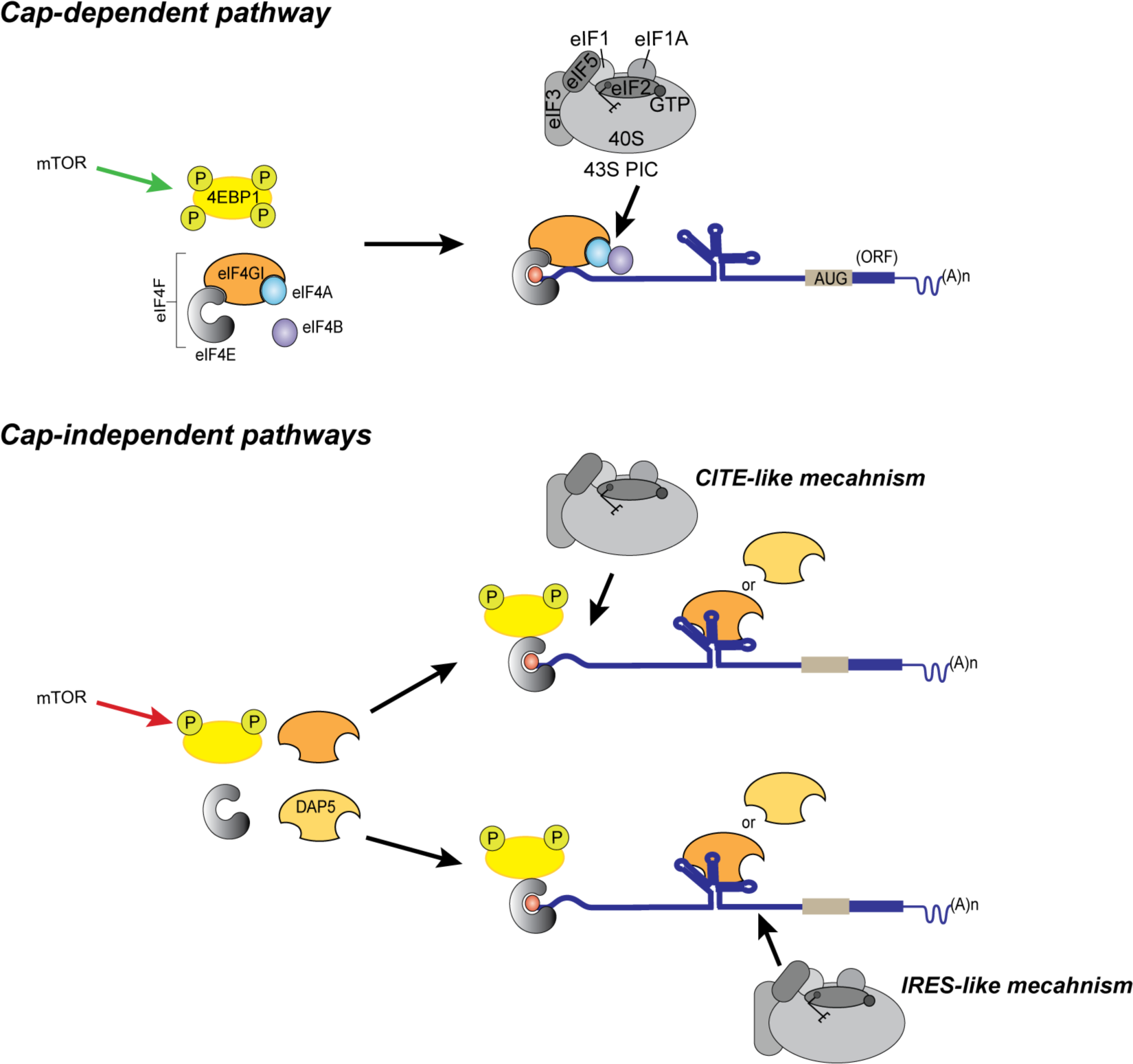
Cartoon summarizing our model for non-canonical, cap-independent translation initiation pathways. The top panel depicts the canonical, cap-dependent pathway, wherein eIF4E binds to the m^7^G-capped mRNA to recruit other eIFs and the 43S PIC. The bottom panel depicts our model for non-canonical, cap-independent pathways, wherein eIF4GI or DAP5 binds to the 5’ UTR and recruits the 43S PIC through either a CITE-like mechanism or an IRES-like mechanism. In all cases, eIF4E interacts with the m^7^G-capped mRNA, but, in the cap-independent pathways, the interaction of eIF4GI with eIF4E is blocked by binding of hypophosphorylated 4EBP1.

The transition from cap-dependent to cap-independent translation of cellular mRNAs under stress conditions has important physiological consequences, particularly in disease states, such as diabetes, cancer, and possibly neurological disorders (7-12). Under stress conditions, the levels of 4EBP1, eIF4GI, and DAP5 are elevated (2,7,8,14,15). Here we have identified key interactions of eIF4GI and DAP5 through which cellular mRNAs containing either IRESs or CITEs may bypass the 4EBP1-mediated depletion of eIF4E and undergo cap-independent initiation and expression. Identification of specific interactions that facilitate the transition from cap-dependent to cap-independent translation promises to facilitate a better understanding of non-canonical translation mechanisms and to inform the potential development of novel therapies that modulate this transition in order to regulate the cellular response to stress conditions.

## Experimental procedures

### Preparation of RNAs for fluorescence anisotropy-based equilibrium binding studies

DNA templates corresponding to the 5’ UTR of HIF-1α (294 nts, GenBank Accession Number: AH006957.2) (55), the 5’ UTR of FGF-9 (177 nts, GenBank Accession Number: AY682094.1) (26), the 5’ UTR for one isoform of p53 (p53_A_) (136 nts, GenBank Accession Number: JN900492.1), the 5’ UTR for a second isoform of p53 (p53_B_) (117 nts, GenBank Accession Number: MG595994.1) (28,45), the ferritin IRE (56), the EMCV J/K IRES (nts 680-786) (31,44), the 101-nucleotide polyUC, and the 5’ UTR of β-actin (84 nts, GenBank Accession Number AK301372.1) (29) were purchased from Integrated DNA Technology (IDT) and the corresponding RNAs were synthesized *via in vitro* transcription using the HiScribeTM T7 Quick High Yield RNA Synthesis Kit (New England Biolabs Inc.) following the manufacturer’s protocol. RNA from transcription reactions were purified using the RNA Clean and Concentrator (RCC) Kit from Zymo Research following the manufacturer’s protocol. Purified RNA transcripts were labeled with fluorescein at their 5’ termini using the 5’ EndTag DNA/RNA Labeling Kit from Vector Laboratories following the manufacturer’s protocol.

### Preparation of eIF4GI_557-1599_, eIF4GI_682-1599_, and DAP5

The plasmids for expression of eIF4GI_557-1599,_ and eIF4GI_682-1599_ were a generous gift from Dr. Christopher Fraser (University of California at Davis). The codon-optimized eIF4GI_682-1599_ construct (34) includes the minimal sequence for IRES-mediated cap-independent translation initiation (31) and is similar in domain structure to full-length DAP5. This construct has an N-terminal, 6x-histidine tag followed by a Flag tag in pET28c vector. We introduced a Tobacco Etch Virus (TEV) protease cleavage site following the Flag tag using the Q5® Site-Directed Mutagenesis Kit from New England Biolabs Inc following the manufacturer’s protocol. The eIF4GI_557-1599_ construct was in fastback vector. It was PCR amplified using a forward primer containing a NcoI site and a 6x-histidine tag followed by a TEV protease site and a reverse primer containing an XhoI site. The amplified PCR product was subcloned in pET28c at the NcoI-XhoI site and used to express and purify the eIF4GI_557-1599_ protein. The plasmid encoding full-length human DAP5 with an N-terminal 6x-histidine tag was purchased from Genscript (Piscataway, NJ). All the proteins were recombinantly expressed in *E. coli BL*21-CodonPlus (DE3)-RIL cells (Agilent) and were purified using a combination of Ni^2+^-nitro-tri-acetic acid (Ni-NTA) affinity and heparin affinity columns, as previously described (25,34). Briefly, the proteins were first purified from bacterial cell lysates using His-Trap HP (Ni-NTA) columns (GE Healthcare Life Sciences), as per the manufacturer’s instructions. The purified 6x-histidine tagged proteins were dialyzed overnight against Storage Buffer (20 mM HEPES pH 7.6, 200 mM KCl, 10 mM β-mercaptoethanol, 10% glycerol) in the presence of TEV protease to cleave off the tags. The untagged proteins were further purified and concentrated using 1 mL HiTrapTM Heparin HP columns (GE Healthcare Life Sciences). The eluted proteins were analyzed on 10% SDS-PAGE gels and pure fractions (>95% purity) were pooled and dialyzed overnight against Storage Buffer. The concentrations of the purified, concentrated proteins were quantified using Coomassie Protein Assay Reagent (Thermo Scientific) and were aliquoted and stored at -80°C.

### Fluorescence anisotropy-based equilibrium binding assays

Fluorescein-labeled RNAs were diluted to 100 nM using Folding Buffer (20 mM HEPES pH 7.5 and 100 mM KCl) and heated to 90 °C for 2 min and slowly cooled over 1 hr to room temperature. MgCl_2_ was then added to the solution to a final concentration of 1 mM, the solution was gently mixed and incubated on ice for about 1 hr. Fluorescence anisotropy measurements for assessing the binding of eIF4GI constructs or DAP5 to the fluorescein-labeled RNAs were performed using the equilibrium titration module of an SF-300X stopped-flow fluorimeter (KinTek Corporation, Austin, TX). Fluorescein-labeled RNAs were excited at 495 nm and emission was detected using a 515 nm high-pass filter (Semrock, Rochester, NY). Equilibrium binding titrations began with a 200 μL sample of 100 nM fluorescein-labeled RNA in the titration buffer (20 mM HEPES, pH 7.6, 100 mM KCl, and 1 mM MgCl_2_) and 20-50 data points were collected for each anisotropy measurement by automated continuous injection of 20 μL of 10 μM eIF4GI_557-1599_, eIF4GI_682-1599_ or DAP5 over a period of 30 min at a temperature of 25 °C. Note that the first reading is taken in the absence of protein. Using the Origin 2018b software package, the data were fitted to a nonlinear, single-site, equilibrium binding equation of the form:

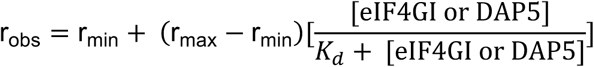

where r_obs_ is the observed anisotropy value, r_min_ is the minimum anisotropy value in the absence of eIF4GI or DAP5, r_max_ is the final saturated anisotropy value, [eIF4GI or DAP5] is the concentration of eIF4GI_557-1599_, eIF4GI_682-1599_, or DAP5, and *K*_d_ is the equilibrium dissociation constant. The chi-squared values (χ^2^) that represented the statistical goodness of fit were always close to 1 and are reported in Table 1. Fitting data to a two-site model did not improve the fit as judged by (χ^2^) values. The equilibrium binding titration of each 5’ UTR was performed three times and fit independently for *K*_d_. The fitted *K*_d_s were then averaged and the standard deviations were calculated (Table 1).

### Preparation of UTR-Luc reporter mRNAs for luciferase-based gene expression reporter assays

The UTR-Luc mRNA constructs for the luciferase gene expression reporter assays were generated from the BlucB plasmid (57), which contains a firefly luciferase gene flanked by 5’- and 3’ UTR sequences of the Barley Yellow Dwarf Virus (BYDV) genomic RNA. To generate each UTR-Luc mRNA reporter construct, a sequence containing the T7 promoter followed by the target 5’ UTR was cloned into the BlucB plasmid vector upstream of the firefly luciferase coding region, after removing the BYDV 5’ UTR. Briefly, all UTR-Luc reporter constructs were PCR amplified with the forward primers containing a NotI site and reverse primers containing a BssHII shown in Table 1 and then digested with NotI and BssHII. All the restriction enzymes were purchased from New England Biolabs (NEB) and restriction digests were performed according to the manufacturer’s instructions. The PCR-amplified sequences were ligated into a NotI- and BssHII-digested BlucB plasmid using DNA ligase (NEB) and transformed into *E. coli* DH5-α competent cells. Five colonies were selected, grown overnight in Luria-Bertani (LB) growth media supplemented with 100 μg/mL ampicillin, and used to isolate plasmid DNA using the QIAprep® Spin Miniprep kit from Qiagen and following the manufacturer’s protocol. To assess whether the availability of a 5’ end is required for the translation of these reporter mRNAs, a highly stable stem-loop (36) was inserted at the 5’ end of each UTR-Luc reporter (hp-UTR-Luc mRNA) using site-directed mutagenesis with the primers in Table 1. All clones were confirmed by sequencing (Genewiz). To generate a linearized plasmid DNA template for *in vitro* transcription, plasmid DNAs were linearized using KpnI so as to remove the 3’ UTR from the UTR-Luc mRNA reporter construct. The resulting linearized DNA was purified using the GeneJET gel extraction and DNA cleanup Micro Kit from GeneJET as per the manufacturer’s instructions. DNA templates were *in vitro* transcribed using T7 RiboMax Large Scale RNA Production Kit (Promega) following the manufacturer’s protocol. ApppG (NEB) or Ribo m^7^GpppA Cap Analog (Promega) were added to the transcription mix in an ApppG or Ribo m^7^GpppA : GTP ratio of 10 : 1 to get mRNA transcripts with non-functional and functional caps, respectively. Capped RNAs were poly A tailed (pA) using the Poly (A) Tailing Kit (Invitrogen) following the manufacturer’s protocol. The resulting capped and polyadenylated mRNAs were then purified using RNA Clean and Concentrator (RCC) Kit (Zymo) following the manufacturer’s protocol.

### Western Blots and depletion of eIF4GI and DAP5 from the rabbit reticulocyte lysate

Monoclonal mouse primary antibodies for DAP5, eIF4GI, eIF4E, eIF2β and eIF4AI/II (Santa Cruz Antibodies) were used for Western blot and/or immunoprecipitation experiments. For the Western blot experiments, the specific antibodies were diluted to 1 : 1000 in PBST (Phosphate-Buffered Saline-Tween) buffer containing 1 % BSA. The blots were incubated overnight in the primary antibodies at 4 °C with constant shaking. The membranes were washed three times with PBST and then incubated with horseradish peroxidase (HRP)-goat anti-mouse secondary antibody (1 : 5000 dilution, Invitrogen) for 1 hr at room temperature. After three subsequent washes in PBST, the membrane was developed using enhanced chemiluminescent substrate (SuperSignal™ West Femto, Thermoscientific). For the depletion of eIF4GI and DAP5 from the RRL, 20 μL of either eIF4GI or DAP5 capture antibodies were incubated with PureProteome™ Protein A/G mix magnetic Beads (EMD Millipore Corporation) at room temperature with continuous mixing for 1 hr. The bead-antibody complexes were washed for 10 secs with 500 μL PBS. The bead-antibody complexes were captured using a magnetic stand (Promega) and the suspensions were removed. The washing step was repeated 2 more times with PBS and then with the wash buffer (20 mM HEPES-KOH, pH 7.5, 50 mM KCl, 75 mM KOAc, 1 mM MgCl_2_). The nuclease-treated RRLs (Promega) were incubated with these preformed bead-antibody complexes at 4°C with continuous shaking for 1 hr. The magnet was re-engaged to capture the bead-antibody-protein complex and the resulting depleted RRL (RRL_(–)4GI_ or RRL_(–)DAP5_) was collected and used for translation. To determine the extent of depletion, Western blot assays were performed after resolving the samples on a 4-15 % Tris-HCl gradient gel (Bio-Rad Laboratories).

### Luciferase-based gene expression reporter assays

Gene expression was achieved by translating the UTR-Luc mRNAs *in vitro*, using the nuclease treated RRL *in vitro* translation system from Promega. The RRL was made more cap-dependent by addition of 75 mM KCl (58). Each 25 μL reaction contained 70 % v/v of RRL, RRL_(–)4GI_ or RRL_(–)DAP5_ (Promega) supplemented with 0.5 mM MgCl_2_, 0.02 mM amino acid mixture, 10 U/μL RiboLock RNase Inhibitor (Thermoscientific), and varying concentrations of purified eIF4GI constructs, or DAP5 as indicated in the figure legends. 4EGI-1 chemical (Selleck Chemicals) was added to the translation mix when indicated at a concentration of 0.2 mM. Briefly, 1 μg of UTR-Luc mRNA was added to the RRL, RRL_(–)4GI_ or RRL_(–)DAP5_ *in vitro* translation mixture that had been pre-incubated at 30 °C for 10 min following the addition of the specified concentration of eIF4GI constructs or DAP5. The resulting *in vitro* translation reaction was then incubated at 30 °C for 1 hr and stopped by the addition of 60 μM puromycin. Firefly luciferase activities were then assayed using a Glomax 96 microplate illuminometer (Promega). To achieve this, 3 μL of translation reaction was added to 30 μL Bright-Glo Luciferase assay reagent (Promega) and the resulting luminescence was measured in the illuminometer over a spectral wavelength of 350–650 nm and an integration time of 10 s at room temperature. After subtracting the background, measured using an in vitro translation reaction to which no UTR-Luc mRNA had been added, the luminescence data were analyzed and plotted using the Prism 8 software package. Each experiment was repeated three times, and the mean and standard deviation of the luminescence data were reported.

### Statistical analyses

All data were analyzed using student t-test and p values were calculated.

### Data availability

All data are present in the manuscript.

## Acknowledgments

The authors wish to thank Paul Powell for helpful discussions. We thank Prof. Christopher S. Fraser (University of California, Davis) for the recombinant eIF4GI clones. We thank Maya Ramachandran for carrying out some of the preliminary binding studies. The Research reported in this publication was supported by Susan G. Komen for the Cure Postdoctoral Fellowship [PDF12231199 to S.M.]; the National Institute of General Medical Science of the National Institute of Health under Award Numbers; [R01 GM 084288 to R.L.G]; [R01 GM 128239 to D.J.G] and National Center for Advancing Translational Sciences [1UL1TR002384-01 Seed Project to D.J.G]. The content is solely the responsibility of the authors and does not necessarily represent the official views of National Institutes of Health.

## Conflict of interest statement

The authors declare that they have no conflicts of interest with the contents of this article

## SUPPLEMENTARY INFORMATION

**Table S1.** Parameters describing the equilibrium binding of eIF4GI mutants and DAP5 to 5’ UTRs

| 5' UTRs | eIF4GI<sub>557-1599</sub> |  |  | eIF4GI<sub>682-1599</sub> |  |  | DAP5 |  |  |
| --- | --- | --- | --- | --- | --- | --- | --- | --- | --- |
| | $K_d \pm \text{S.D.}$<br>(nM)<sup>a</sup> | Amp.<br>($r_{\text{max}}-r_{\text{min}}$)<sup>b</sup> | $\chi^2$<sup>(c)</sup> | $K_d \pm \text{S.D.}$<br>(nM) | Amp.<br>($r_{\text{max}}-r_{\text{min}}$) | $\chi^2$ | $K_d \pm \text{S.D.}$<br>(nM) | Amp.<br>($r_{\text{max}}-r_{\text{min}}$) | $\chi^2$ |
| EMCV J/K IRES | 63 ± 3.0 | 0.024 | 0.99<br>6 | 175 ± 4.0 | 0.031 | 0.998 | 519 ± 21.0 | 0.045 | 0.988 |
| Ferritin IRE | 15 ± 2.0 | 0.040 | 0.99<br>9 | 18 ± 2.0 | 0.076 | 0.997 | 35 ± 3.0 | 0.069 | 0.998 |
| polyUC | N.A | 0.011 | 0.94<br>6 | N.A | 0.003 | 0.994 | N.A | 0.010 | 0.999 |
| β-actin | N.A | 0.007 | 0.99<br>8 | N.A | 0.006 | 0.911 | N.A | 0.039 | 0.998 |

<sup>a</sup> $K_d$ is the equilibrium dissociation constant<sup>b</sup> $r_{\text{max}}-r_{\text{min}}$ is the amplitude which indicates change in anisotropy<sup>c</sup> $\chi^2$ is the goodness of fit

**Supplementary Figure. 1.**
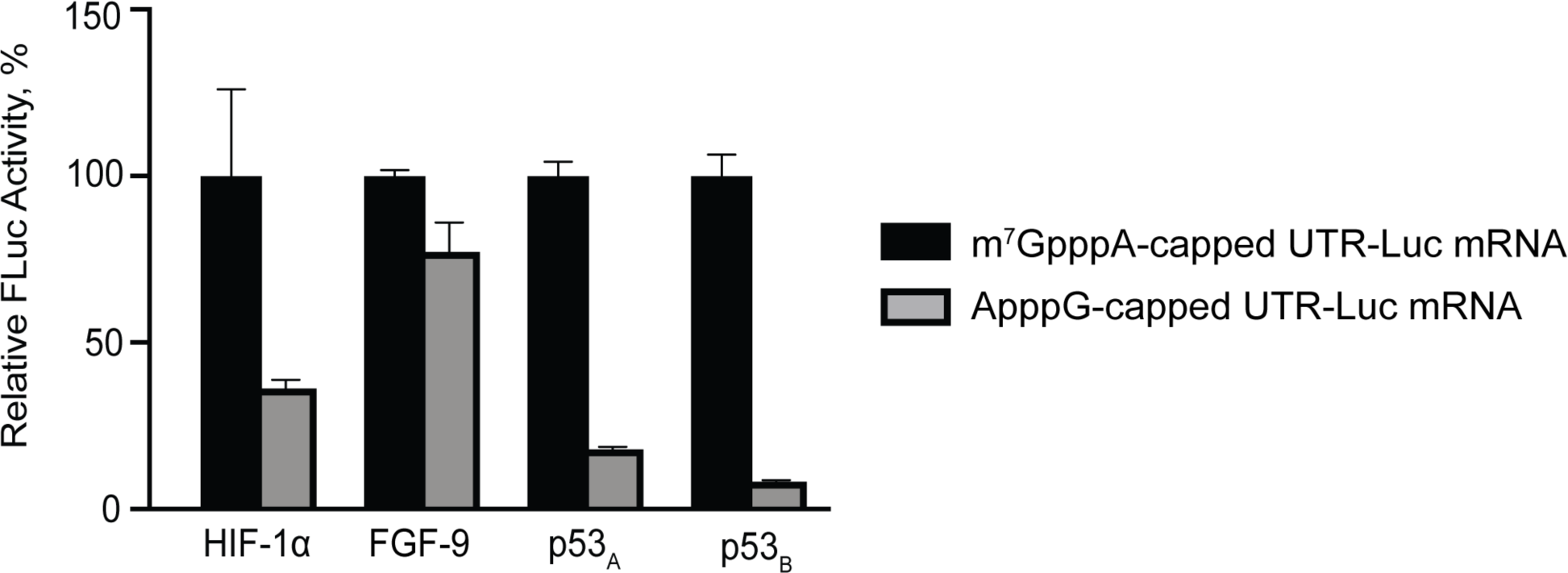
Comparison of cap-dependent and cap-independent translation activities of HIF-1α, FGF-9, p53_A_, p53_B_, and β-actin UTR-Luc mRNAs in RRL. The translation output from the m^7^GpppA-capped UTR-Luc mRNA for each individual mRNA was set to 100% and used to normalize translation levels of ApppG-capped UTR-Luc mRNA.

**Supplementary Figure. 2.**
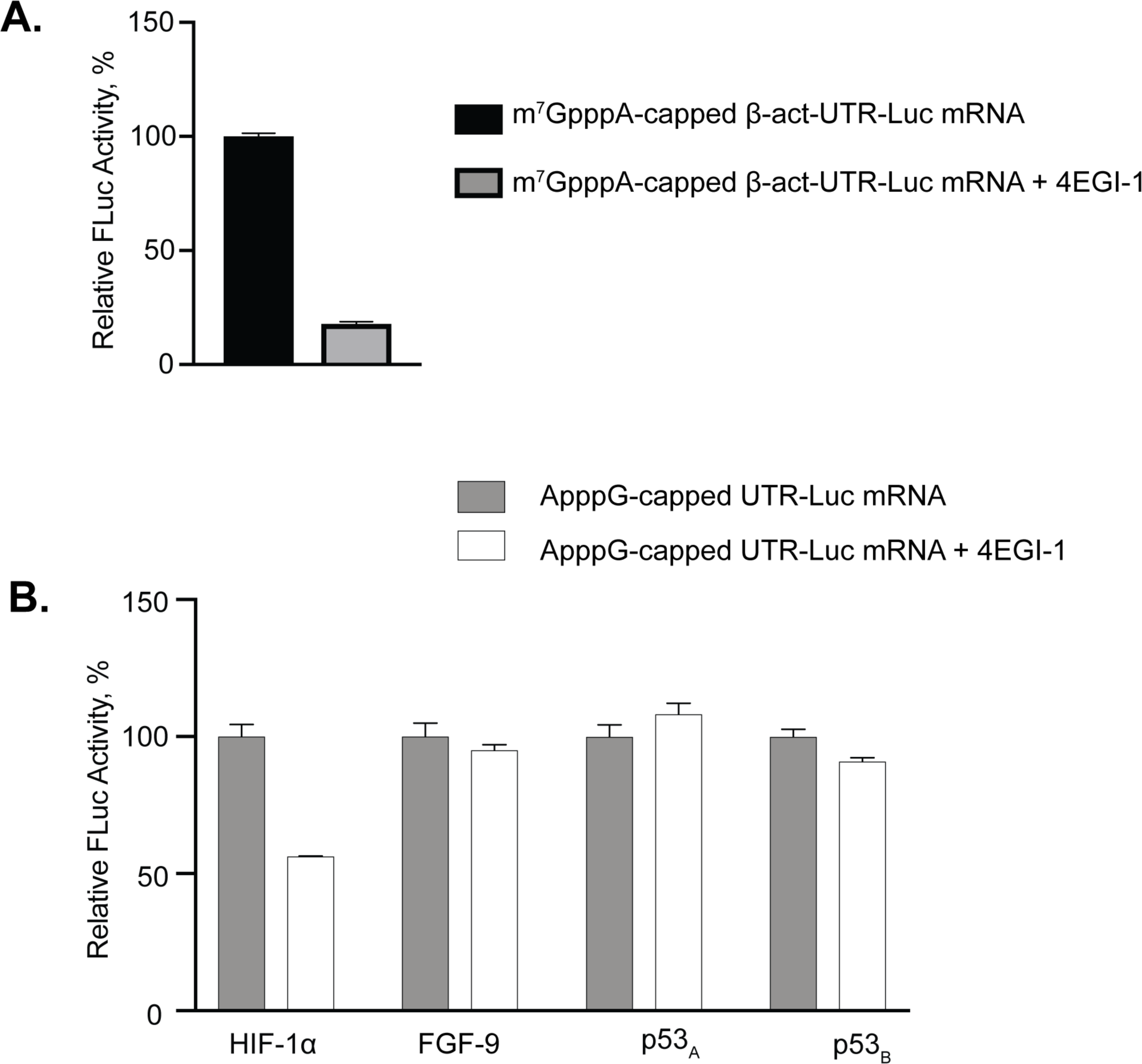
Effect of 4EGI-1 peptide on the translation yields of UTR-Luc mRNAs of **A**. m^7^GpppA-capped β-act-UTR-Luc mRNA. **B**. ApppG-capped UTR-Luc mRNA of HIF-1α, FGF-9, p53_A_, and p53_B_. The RRL was made more cap-dependent by the addition of 75 mM KCl to the lysate.

**Supplementary Figure. 3.**
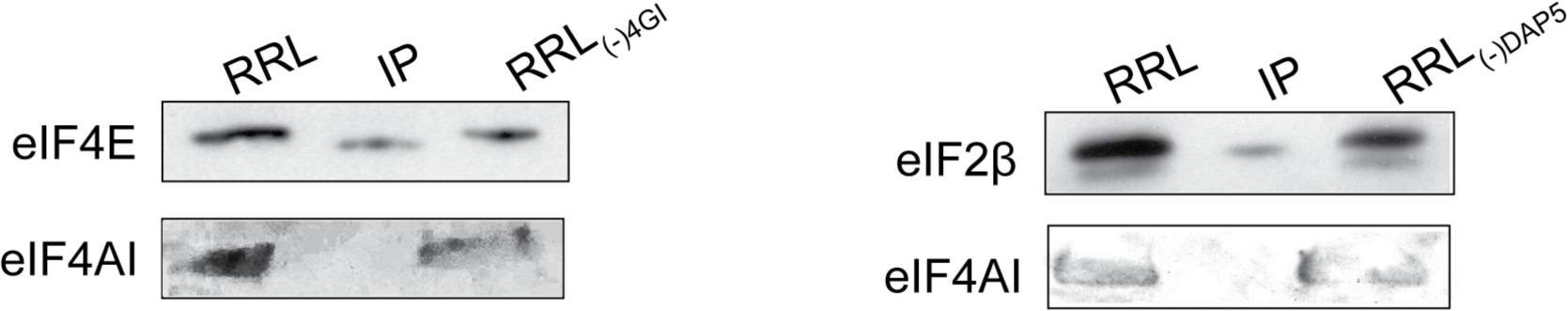
Depletion of eIF4GI or DAP5 have little effect on eIF4E, eIF2β, and eIF4AI levels. IP represents the protein pulled-down by the immunoprecipitation.

## Notes

### Competing Interest Statement

The authors have declared no competing interest.

